# Conversion of mouse-derived hybridomas to Tasmanian devil recombinant IgG antibodies

**DOI:** 10.1101/2023.10.20.563388

**Authors:** Bailey Slyp, Jocelyn M. Darby, Andrew S. Flies

**Author notes:** **CORRESPONDING AUTHORS CONTACT INFORMATION Andrew S. Flies, PhD**, Menzies Institute for Medical Research, College of Health and Medicine University of Tasmania, Private Bag 23, Hobart TAS 7000.

## Abstract

The hybridoma method for production of monoclonal antibodies has been a cornerstone of biomedical research for several decades. Here we convert the monoclonal antibody sequence from mouse-derived hybridomas into a “devilised” recombinant antibody with devil IgG heavy chain and IgK light chain. The chimeric recombinant antibody can be used in functional assays, immunotherapy, and to improve understanding of antibodies and Fc receptors in Tasmanian devils. The process can be readily modified for other species.

## 1 Introduction

Antibodies are an essential tool for the characterization and detection of proteins involved in both normal and pathological functions [1,2]. A wide array of monoclonal antibodies that bind specific proteins are available for humans and select rodent species. Monoclonal antibodies (MAbs) have historically been developed using the hybridoma method of fusing mouse B cells and myeloma cells [3]. The monoclonal antibodies produced using mice (*Mus musculus*) are directly useful for immune phenotyping experiments for the target species but have limited ability for *in vivo* functional assays in species other than mice. This is due to the antibody heavy and light chains being recognised as foreign in other species and the unknown interactions between mouse antibodies and host species Fc receptors. Thus, monoclonal antibodies must be converted to species-matched recombinant antibodies for proper functional assays.

Recombinant monoclonal antibodies (rMAbs) can be produced by cloning the antibody variable region sequences from hybridomas into expression vectors. Recombinant MAbs that retain the mouse variable region but splice it with heavy (e.g., human IgG1) and light (e.g. human kappa) chain constant region antibody genes from another species are considered chimeric rMAbs. After cloning mouse variable regions into an expression vector and validating that the binding to the known target is preserved, the variable regions can be rapidly swapped into alternative expression vectors that encode other heavy chain isotypes (e.g. IgE, IgM) or subclasses (e.g. human Ig2, IgG3, IgG4) [4]. Additionally, DNA expression vectors for rMAbs can easily be shared between organisations without the need for shipping cells on dry ice, as would be needed for hybridomas. The shared expression vectors can be further customised to meeting the needs of research projects.

Immune checkpoint immunotherapy using rMAbs has transformed human oncology in the past decade [5,6]. The monoclonal antibody-based immunotherapies work by manipulating interactions between immune checkpoint protein receptors and ligands and altering survival and expansion of host immune cells. The most potent immune checkpoint immunotherapy targets to date have been CTLA4, PD1, and PDL1. The functional role and binding patterns of these immune checkpoint proteins are unknown for most species due to a lack of species-specific MAbs [7]. Our team has developed and validated anti-PD1 and anti-PDL1 MAb hybridomas for Tasmanian devils (*Sarcophilus harrisii*) to develop potential immunotherapies for transmissible cancers in Tasmanian devils [8].

The original devil facial tumour (DFT1) was first observed in 1996 and demonstrated to be clonally transmissible cancer in 2006 [9]. DFT1 has now spread to most of the island state of Tasmania and has resulted in local population declines of about 80% after the tumour arrives in the area [10]. In 2014 a second, independent devil facial tumour (DFT2) was detected on peninsula in southern Tasmania [11]. Infection by the transmissible DFT1 and DFT2 cells usually results in death of the devil within 3-12 months. No treatments are currently available to treat DFT1 or DFT2.

Here we demonstrate how to convert traditional hybridoma-derived monoclonal antibodies to ‘devilised’ recombinant antibodies (drMAbs) with devil IgG heavy chain and kappa light chain constant regions (**Fig. 1**). We initially assembled and tested a humanised IgG1 anti-6xHis-tag monoclonal antibody expression vector with mouse variable regions and human constant regions (**Fig. 2a-b**). This allowed us to validate the expression vector system using well-characterised variable regions from the anti-His-tag clone 3D5 and created an antibody to serve as a positive and negative control in downstream assays [12]. Next, we amplified the mouse anti-Tasmanian devil PDL1 heavy VDJ chain and light VY chain coding regions from hybridoma clone 1F8 cDNA and assembled this into our human IgG1 expression vector plasmid (**Fig. 2c**) to create a human IgG1 antibody that binds devil PDL1. These recombinant monoclonal antibodies produced were tested for specificity to devil PDL1 by flow cytometry analysis.

**Figure 1.**
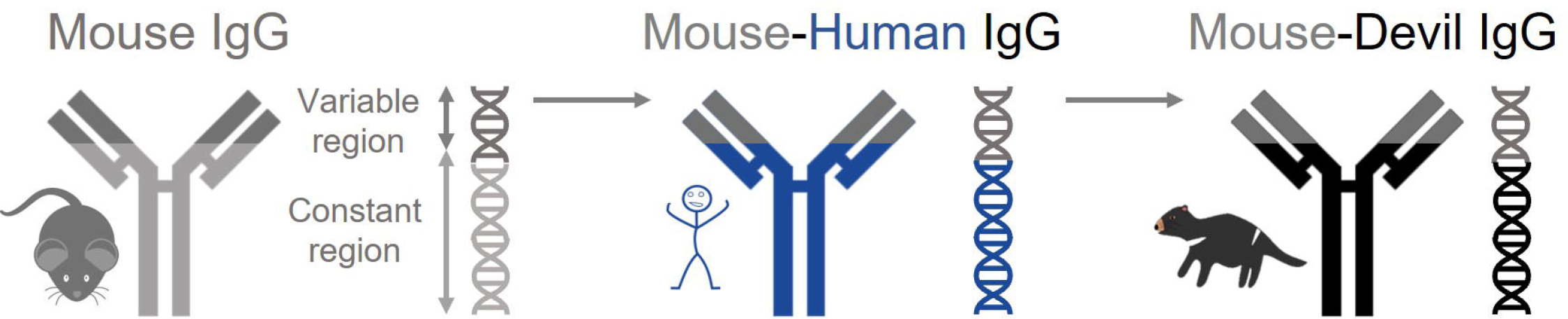
Overview of recombinant antibodies. Antibody variable region DNA is amplified and cloned from the original antibody-secreting cells and transferred into a DNA expression vector for production of the recombinant antibody. Specificity of the antibody is dependent on the variable regions, and thus can be paired with constant regions from other species. The mouse antibody variable region was first combined with the human IgG1 constant region to make a “humanised” mouse-human chimeric antibody. The mouse variable region was then paired with the devil IgG constant region to make “devilised” mouse-devil chimeric antibody.

**Fig. 2.**
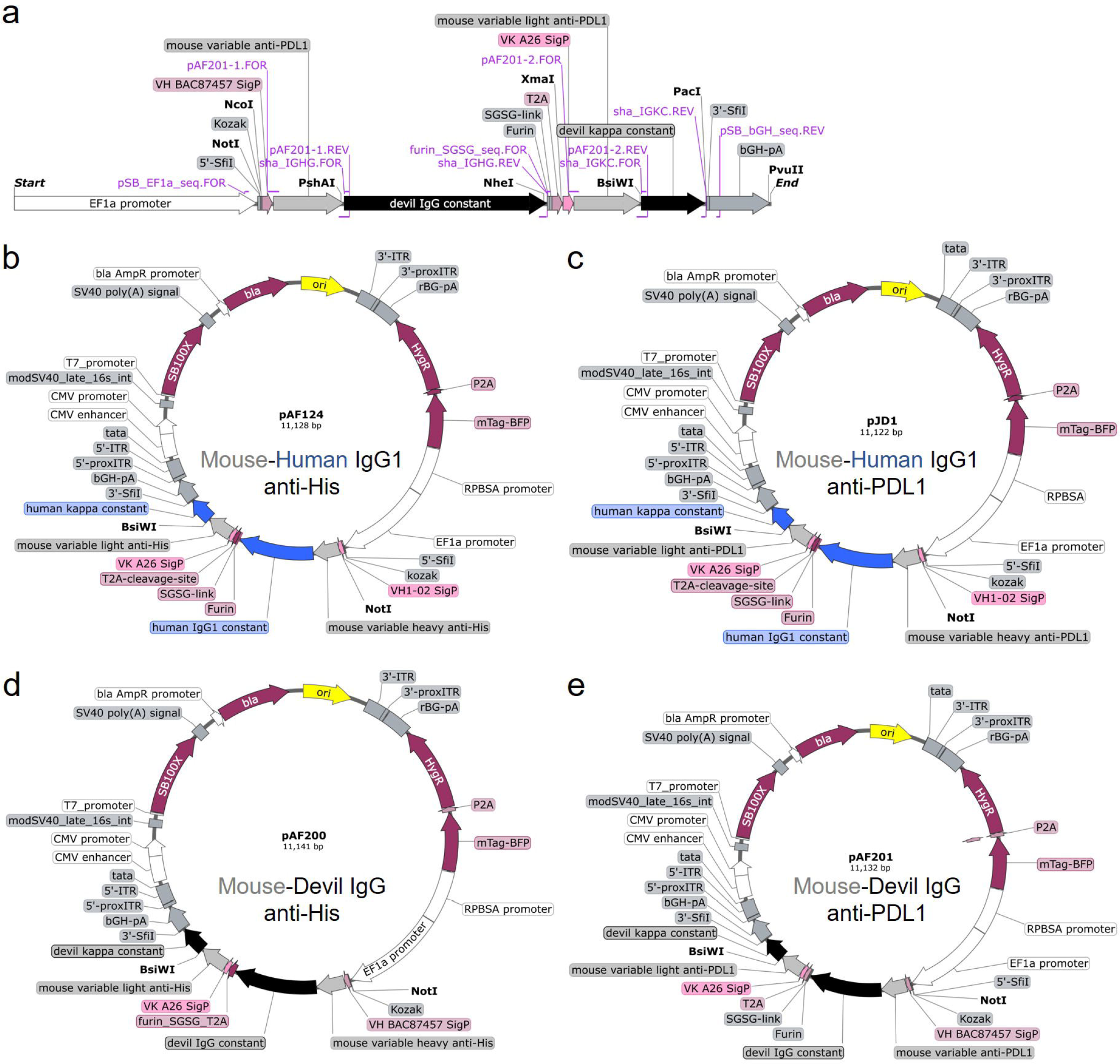
Maps of recombinant antibody expression vectors. (a) Linear map of the human IgG1 anti-6xHis tag antibody expression cassette. The cassette included a human EF1α promotor and bovine growth hormone polyadenylation signal (bGH-pA) to antibody expression. The heavy and light chains were linked by a peptide containing a furin cleavage site and T2A self-cleaving peptide to produce both proteins from a single promotor in a 1:1 ratio. A synthetic RPBSA promotor in the opposite direction was used to co-express mTAG-BFP and a hygromycin resistance gene, linked with a P2A self-cleaving peptide. Full plasmid map for (b) human IgG1 anti-6xHis tag (pAF124), (c) human IgG1 anti-devil PDL1 (pJD1), (d) devil IgG anti-6xHis tag (pAF200), and (e) devil IgG anti-devil PDL1 (pAF201).

After showing that the humanised anti-His and anti-devil PDL1 rMAbs were able to bind their respective protein targets, we began ‘devilising’ the antibodies. We replaced the human IgG1 and kappa sequences with devil IgG and kappa sequences (**Fig. 2d-e**). We produced a Tasmanian devil IgG anti-His antibody first, then amplified and inserted the anti-devil PDL1 variable regions to produce a recombinant monoclonal Tasmanian devil IgG anti-PDL1 antibody. These devilised antibodies were tested by enzyme-linked immunosorbent assay (ELISA) and flow cytometry assays to confirm binding the target proteins. These devilised rMAbs can be used in functional *in vitro* assays using devil immune cells or potentially used for *in vivo* immunotherapy trials for devils with DFT1 or DFT2.

## 2 Materials

### 2.1 Gel electrophoresis

1. 50X TAE buffer: 2 M Tris base, 1 M glacial acetic acid, 50 nM EDTA solution, pH 8.0. Prepare to 1 L using pure water (e.g. Milli-Q water or deionised water) and filter-sterilise the solution through a 0.22 μm membrane.
2. 1X TAE buffer: Dilute 20 mL 50X TAE buffer in 980 mL of pure water.
3. 1% agarose gel: Agarose powder, 1X TAE buffer, Gel Red Nucleic Acid Stain. Dissolve 0.5 g agarose powder in 50 mL of 1X TAE buffer in a microwave for 2-3 minutes with intermittent swirling. Add 5 μL GelRed nucleic acid stain to molten agarose gel and swirl to mix. Pour into a gel casting tray and cool for 10 minutes to set.

### 2.2 Bacterial transformation

1. 80% glycerol solution: Combine 80 mL glycerol with 20 mL pure water. Autoclave for 20 minutes.
2. Luria-Bertani (LB) broth medium with ampicillin: Dissolve 10 g tryptone, 5 g yeast extract, and 10 g NaCl in 1 L ddH_2_O. Autoclave solution at 120 °C for 30 minutes. Once cooled to 55 °C, add 100 mg ampicillin.
3. LB agar with ampicillin: Make up using the LB broth recipe to 980 mL. Add 15 g agar to the ddH_2_O at the same time as the other reagents, then top up with pure water to 1 L. Autoclave and then place media in 55 °C water bath to cool and maintain temperature to avoid having the agar turn solid. Add 100 mg of ampicillin after the media has cooled to 55 °C. Pour LB agar into sterile petri dishes immediately. Allowing the agar to cool and solidify in a biosafety cabinet with the lids slightly open will reduce water condensation on the plates. Store plates at 4 °C with the lids closed in a sealed bag after the agar has fully cooled and solidified.
4. Phosphate-buffered saline (PBS): Dissolve a single PBS tablet (Gibco, USA) 140mM NaCl, 2.68 mM KCl, 10mM NaP, in 500 mL Milli-Q water. Autoclave at 120 °C for 30 minutes.

### 2.3 Cell transfection

1. Complete RPMI with 5% foetal bovine serum (cRF5): RPMI 1640 medium with 2 mM L-glutamine, 5 mL antibiotic antimycotic, 10 mM HEPES, 50 μM 2-mercaptoethanol, 25 mL heat-inactivated FBS. Store at 4 °C protected from light. Heat to 37 °C before using. Keep all cell culture media sterile by working laminar flow biosafety cabinet.
2. Polyethyleneimine (PEI) (1 ug/ mL): Polyethyleneimine, linear. Mix 100 mg Polyethyleneimine, linear in 100 mL Milli-Q water. Cover top of container and heat to 80 °C and stir for three hours, or until solution is mostly clear. Cool to room temperature. Adjust pH to 7 and then sterile filter through 0.22 μm membrane. Aliquot 1 mL into vial and store at -20 °C.

### 2.4 ELISA

1. ELISA coating buffer: Dissolve 0.42 g sodium bicarbonate (Na_2_CO_3_) and 1.8 g sodium carbonate (NaHCO_3_) in 450 mL Milli-Q water. Adjust pH to 9.5 and top up to 500 mL with water.
2. ELISA antibody dilution / blocking buffer: Dissolve 1 g bovine serum albumin and add 0.1 mL of Tween®-20 into 100 mL of PBS using a magnetic stirrer. Store at 4 °C and use within 1 week.
3. Phosphate Buffered Saline with Tween®-20. Add 0.5 mL of Tween®-20 into 1 L of PBS and mix using a magnetic stirrer.

### 2.5 Flow cytometry

1. Flow cytometry buffer: Dissolve 5 g bovine serum albumin in 996 mL PBS using a magnetic stirrer at 4 °C. Add 4 mL of 10% sodium azide solution. Filter-sterilise through a 0.22 μm membrane and store at 4 °C.
2. Optional: 10% sodium azide solution (hazardous). Wear a mask, gloves, and safety glasses when working with sodium azide (NaN_3_). Dissolve 10 g of sodium azide in 100 mL of pure water.

## 3 Methods

### 3.1 Recombinant antibody expression vectors

The expression vector plasmid is essential for the successful transfection of cell lines and production of recombinant antibodies (*see* **Note 1**). We used our previously validated all-in-one sleeping beauty transposon plasmid system as base vectors for this study [13,14]. The all-in-one sleeping beauty vectors contain the sleeping beauty transpose derived from the pCMV(CAT)T7-SB100 (<u>Addgene # 34879</u>) and the sleeping beauty transposon cassette (<u>Addgene # 60515</u>) [15,16]. Transfection of mammalian cells with these plasmids and subsequent drug selection yields insertions of the transposon cassette that are stable for at least three months even in the absence of ongoing drug selection (*see* **Note 2**) [17].

We modified the all-in-one sleeping beauty vectors to produce the base recombinant antibody expression vector (pAF125) that was used for downstream cloning. The human IgG1 and kappa light chain constant regions were derived from the plasmid pVITRO1-dV-IgG1/κ (<u>Addgene # 52213</u>) [18]. The IgG1 and kappa chains in human IgG1 vectors were linked by a Furin-SGSG-T2A linker. The production of high-quality monoclonal antibodies relies on balancing the expression of the light and heavy chain antibody subunits. Linking the IgG1 and kappa sequences with the self-cleaving peptide T2A enables co-expression at equal amounts in one open reading frame [19]. Furin enables removal of the T2A sequence from the heavy chain after self-cleavage [19].

The pAF125 plasmid is an empty recombinant antibody vector for human IgG1 and kappa without variable regions. Mouse variable regions that bind 6xHis tags were inserted into the pAF125 plasmid to make the pAF124 plasmid (**Fig. 2b**). Both plasmids are available from Addgene (reference numbers pending). The pAF124 plasmid that encodes a recombinant human IgG1 antibody that binds 6xHis tags is the starting point for this protocol. The protocol should work for most other mouse hybridoma cells. We used anti-devil PDL1 clone 1F8 to illustrate the process here.

### 3.2 Assembly of human IgG1 anti-devil PDL1 (pJD1) expression vector

#### 3.2.1 Isolate RNA from hybridoma for devil PDL1 and make cDNA

1. Culture monoclonal hybridoma to obtain at least 1 million viable cells.
2. Extract RNA from a hybridoma clone producing mouse monoclonal anti-devil PDL1 antibodies using a Nucleospin® Plus RNA extraction kit, according to the manufacturer’s instructions.
3. Remove extra DNA with a TURBO DNA-*free*™ Kit (ThermoFisher Scientific, USA), according to the manufacturer’s instructions.
4. Synthesize cDNA from the extracted hybridoma RNA using a GoScript™ Reverse Transcription System 100X reaction kit (Promega, USA), according to the manufacturer’s instructions.

#### 3.2.2 PCR amplify mouse anti-devil PDL1 variable regions

1. Degenerate primers to amplify mouse heavy and light chain variable regions [4] were modified to add 16-20 bp to the 5’ end to create extensions that overlap the pAF124 vector. Mouse variable heavy (mVH) and mouse variable light kappa (mVLk) primers with extensions are shown in Table 1. The primers were mixed at the ratios shown in Table 1 to form a 100uM primer cocktail that covers the most common mouse variable region genes.
2. Add 10 μL of Q5® Hot Start High-Fidelity 2X Master mix (NEB), 1 μL of cDNA, and 3 μM of the forward and reverse primer cocktails. The total volume will be 20 μL.
3. Touchdown PCR that initially decreases the annealing temperature by 1 °C per cycle for the first ten cycles before returning to a higher annealing temperature for subsequent cycles is used with the following cycling conditions. Denature the DNA at 98 °C for 1 minute, 10 cycles of 98 °C for 10 seconds, 65-55 °C for 15 seconds (decreasing by 1 °C per round), and 72 °C for 45 seconds. This is immediately followed by 30 cycles of 98 °C for 10 seconds, 65 °C for 15 seconds, and 72 °C for 45 seconds, followed by a final extension at 72 °C for 2 minutes.
4. The linker fragment from the pAF124 plasmid is amplified from the 5’ end of the IgG1 constant chain to the 3’ end of the light chain signal peptide using forward primer Linear_IgG1-H_FOR and Linear_Krev. Although not optimised, this reaction can be run using the same conditions above.
5. Pour 50 mL of 1% agarose gel into a gel casting tray and comb to form wells. Leave for at least 30 minutes to cool and set. Remove the comb and place the gel in a gel box (Bio-Rad, USA). Cover the gel with 1X TAE buffer.
6. Add 4 μL of 6X Gel Loading Dye (New England BioLabs, USA) to each PCR tube. Load 5 μL of DNA ladder into the first lane of the gel.
7. Load 10 μL of the PCR solutions into the other wells.
8. Run the gel at 100 V for 40-60 minutes.
9. Briefly image under blue light and cut out the bands containing the variable regions and the linker (*see* **Note 3**).
10. Purify the DNA using the NucleoSpin® PCR and Gel Clean Up Kit (Macherey-Nagel, USA), according to the manufacturer’s instructions.
11. Quantify the DNA concentrations using a Nanodrop or other spectrophotometer.
12. Re-amplify the variable region fragments using the appropriate nested primers.
  a. VH1-02_nested.FOR and VH1-02_nested.REV for the heavy chain variable region
  b. VK-A26_nested.FOR and VK-A26_nested.REV under the same cycling conditions.
13. Purify the DNA using the NucleoSpin® PCR and Gel Clean Up Kit (Macherey-Nagel, USA), according to the manufacturer’s instructions.
14. Quantify the DNA concentrations using a Nanodrop or other spectrophotometer.

**Table 1.**
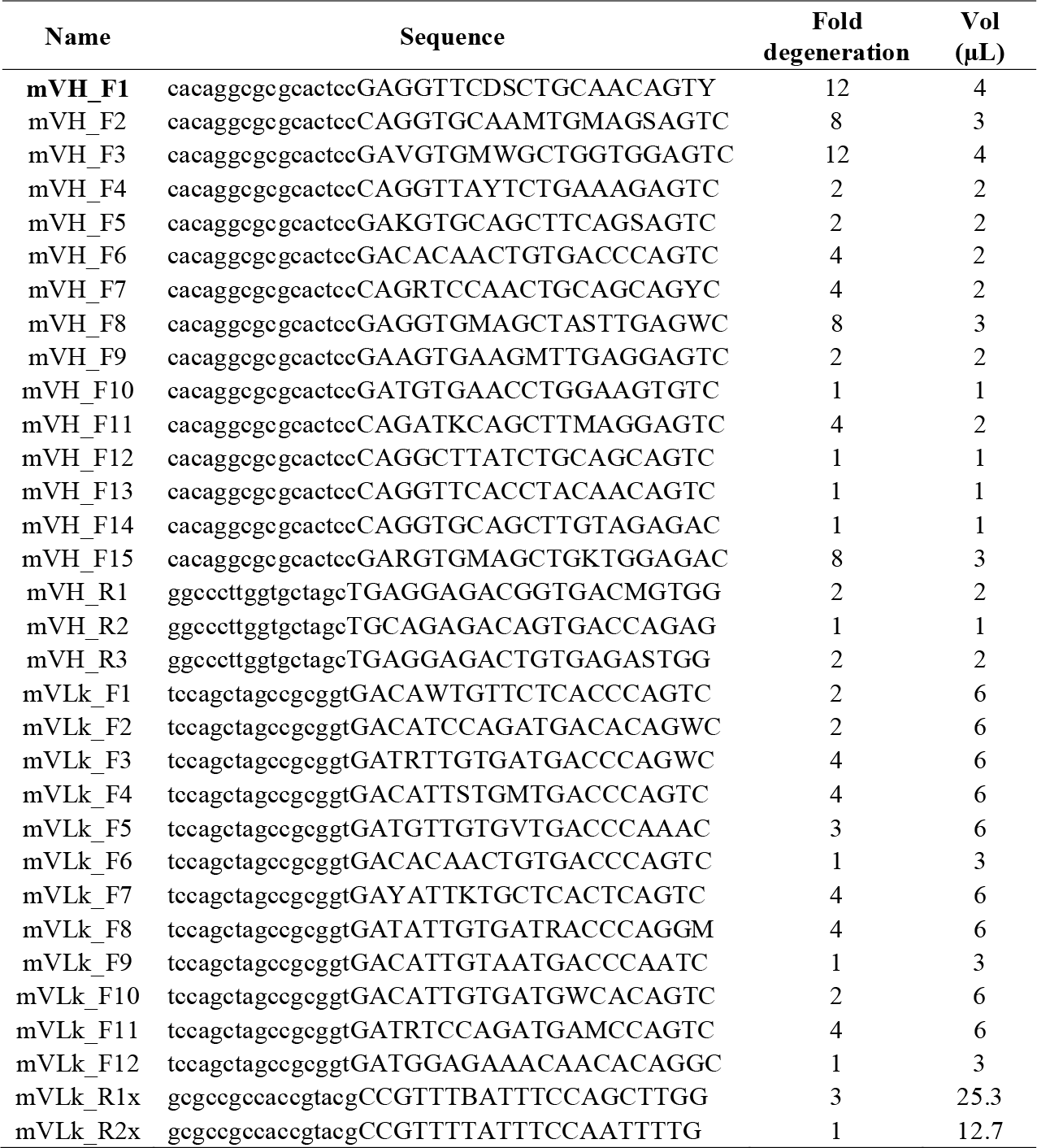
Mouse anti-devil PDL1 variable region primer sequences.

#### 3.2.3 Restriction digest of the pAF124 human IgG1 anti-His expression vector

1. In a 50 μL reaction add 5 μL of 10X Cutsmart® buffer (New England Biolabs, USA), 3 μg of expression vector DNA, 3 μL of NotI-HF (NEB) and 3 μL of BsiWI-HF (NEB), and nuclease-free water. Incubate overnight at room temperature.
2. Incubate the solution at 37 °C for 15 minutes, 65 °C for 20 minutes, then cool to 4 °C (*see* **Note 4**).
3. Dephosphorylate the vector by adding 6 μL of Antarctic phosphatase buffer (NEB) and 3 μL of Antarctic phosphatase. Incubate at 37 °C for 1 hour at room temperature then heat inactivate at 80 °C for 2 minutes.
4. Set up a 1% agarose gel as previously described (section 3.2.2) (*see* **Note 5**).
5. Run the gel at 100 V for 1 hour. Briefly image under blue light and cut out the band containing the digested vector.
6. Purify the vector DNA using the NucleoSpin® PCR and Gel Clean Up Kit (Macherey-Nagel, USA), according to the manufacturer’s instructions.

#### 3.2.4 Use Gibson assembly to add mouse anti-devil PDL1 variable regions to a human IgG1

1. Dilute the digested human IgG1 expression vector to a concentration of 5 ng/μL.
2. On ice, add nuclease-free water, 5 μL of 2X NEBuilder® master mix, and the digested vector and PCR amplicons at a 1:3 molar ratio to a total of 10 μL in a sterile PCR tube. Incubate at 50 °C for 1 hour. Alongside this reaction also include a positive and negative control (*see* **Note 6**).
3. Retrieve NEB DH5a cells from cryostorage and thaw on ice (*see* **Note 7**).
4. Gently pipette 2 μL of assembled recombinant plasmid vector product into 25 μL of thawed DH5a bacterial cells 5 times and gently tap the tube 5X to mix DNA and bacteria (*see* **Note 8**).
5. Incubate cells on ice for 30 minutes, then heat shock the bacteria by holding the tube in a 42 °C water bath for 30 seconds, then immediately place the tubes ice for 2 minutes (*see* **Note 9**).
6. Add 950 μL of SOC media to the bacteria in the tube and incubate at 37 °C for 60 minutes on an orbital shaker at 200 RPM with the tubes at a 45-to-60-degree angle.
7. Spin tubes at 8000 x g for 30 seconds. Discard media, leaving 150 μL in which to resuspend the transformed bacterial cells. Spread 100 μL of the transformed bacteria onto LB agar plates containing 100 μg/mL ampicillin using a cell-spreader and culture overnight at 37 °C with the lid on the bottom and the agar on top (*see* **Notes 10-11**). Store the additional 50 μL at 4 °C as backup. Remember to discard later.
8. Observe the agar plates for bacterial growth and record the number of colonies (*see* **Note 12**). Select 5 single colonies from the recombinant plasmid vector agar plate using sterile pipette tips and inoculate each separately into 10 μL of PBS and store at 4 °C.
9. In a sterile PCR tube, add 7.8 μL of nuclease-free water, 10 μL of OneTaq® Hot Start Quick-Load 2X Master Mix (NEB biolabs, USA), 0.6 μL of pSB_EF1a_seq.FOR and pSB_bGH_seq.REV primers, and 1 μL of bacteria-PBS solution (*see* **Note 13**). Use a base vector plasmid (pAF124) as a positive control and an irrelevant plasmid or water only as the negative control.
10. PCR the solutions in a Thermal Cycler under the following cycling conditions – 94 °C for 3 minutes, 25-35 cycles of 94 °C for 15 seconds, 60 °C for 15 seconds, and 68 °C for 1 min/ kB of amplicon length (e.g. 2 minutes and 30 seconds), followed by a final extension of 68 °C for 5 minutes. Run a negative and positive control alongside the colony PCR lanes in the gel.
11. Setup and run a 1% agarose gel as previously described.
12. Check the amplicon sizes to determine if they are approximately the correct size based on the plasmid design.
13. For clones that have the correct sized amplicon, retrieve the PBS containing the original bacterial colony and transfer 5 μL into 10 mL of LB broth containing 100 μg/ mL ampicillin in a 50 mL tube. Incubate overnight at 37 °C with the tube at a 45 ° angle at 200 RPM.
14. Combine 750 μL of amplified bacterial culture and 250 μL of 80% glycerol solution (section 2.2.1) in a 1 mL cryovial and store at -80 °C for future use.
15. Pellet 3-5 mL of bacterial broth at 3000 x g for 10 minutes and discard the supernatant.
16. Isolate the assembled recombinant vector from the transformed bacteria using the Nucleospin® Plasmid EasyPure Kit (Macherey-Nagel, USA) according to the manufacturer’s instructions (*see* **Note 14**).
17. Sequence the full insert from 3’ end of the EF1a promotor to the 5’ end of the light chain constant region using the pSB_EF1a_seq.FOR and pSB_bGH_seq.REV primers. Primers hsa_IGHG1_seq-1.REV, hsa_IGHG1_seq-1.FOR, furin_SGSG_seq.FOR, and hsa_IGHG1_seq-2.REV should provide full coverage of the region of interest. We have omitted the sequencing details here, as commercial sequencing facilities are available in many locations.

### 3.2.5 Transfect CHO cells with human IgG anti-devil PDL1 expression vector

Chinese Hamster Ovary (CHO) cells were maintained in a monolayer in T75 flasks with 10 mL of cRF5 and incubated at 37 °C with 5% CO_2_. Cells were passaged in biosafety cabinets under sterile conditions once they reached ∼80% confluency. Passaging involved the removal of the supernatant from the cell cultures and washing with 3 mL sterile PBS. Three mL TrypLE express enzyme (1x) was added to the flask and incubated for 5 minutes at 37 °C with 5% CO_2_ to enzymatically detach the adherent cells from the flask. Cells were mechanically agitated by gentle tapping of the flask. Two mL of cRF5 was added and pipetted along the base of the flask to ensure detachment and collection of all cells. The cell solution was transferred to a 50 mL tube and centrifuged at 200 g for 4 minutes. Supernatant was discarded and cells resuspended in cRF5 for counting using a haemocytometer. The desired number of cells were then transferred to a flask for continued cell culture or into another vessel for downstream analysis.

1. Seed a 6-well plate with 2.5x10^5^ CHO cells in 2.5 mL of cRF5 per well. Incubate overnight at 37 °C 5% CO_2_.
2. Cell should be 50-70% confluent prior to transfection.
3. Thaw PEI transfection reagent on ice.
4. For each well to be transfected – dilute 2 μg of recombinant human IgG1 anti-His expression vector plasmid DNA in PBS to a volume of 100 μL. In a separate tube dilute 6 μg PEI (3:1 PEI:DNA ratio) in 100 μL of PBS.
5. Add the PBS:PEI mix to the DNA. Mix by pipetting once. Do not vortex.
6. Incubate at room temperature for 15 minutes.
7. Aspirate the media from the CHO cells. Replace with 2 mL of fresh, warm cRF5.
8. Add 200 μL of the DNA:PEI solution dropwise to the appropriate well.
9. Gently rock the plate to spread the solution throughout the well. Do not swirl.
10. Incubate cells for 4 hours at 37 °C, 5% CO_2_.
11. Replace the cell media with 2 mL of fresh, warm cRF5. Incubate for 24 hours. This step can be skipped if toxicity from the PEI:DNA mix is low.
12. Examine the cells using fluorescent microscopy to estimate the transfection efficiency.
13. Once fluorescent reporter protein is visible, add 800 μg/ mL hygromycin for selection of transformed cells (*see* **Note 15**).
14. Harvest the cell supernatant 8-14 days post transfection and discard the cells (*see* **Note 16**).

#### 3.2.6 Test transfected CHO cell supernatant by flow cytometry to confirm antibody binding

1. Harvest CHO cells stably expressing full-length Tasmanian devil PDL1 (pAF146).
2. Add 2x10^5^ cells to each well of a U-bottom 96-well plate.
3. Centrifuge plate at 300 x g for 5 minutes and discard cell supernatant.
4. Add 100 μL of neat human IgG anti-devil PDL1 CHO cell supernatant to the cells and incubate for 30 minutes on ice. Wash twice with 100 μL of flow cytometry buffer.
5. Add 100 μL of anti-human IgG (Alexa Fluor™ 488) diluted 1 μg in 1 mL flow cytometry buffer. Incubate for 30 minutes on ice. Wash twice with 100 μL flow cytometry buffer.
6. Resuspend cells in 400 μL DAPI in flow cytometry buffer to distinguish live and dead cells.
7. Analyse cells by flow cytometry along with controls (*see* **Note 17**).

### 3.3 Tasmanian devil IgG anti-His (pAF200) and anti-PDL1 (pAF201) assembly

To ‘devilise’ these recombinant antibodies, two g-blocks coding for the Tasmanian devil IgG heavy and light chain constant regions (IGHC & IGKC) and furin-GSG-T2A linker were obtained from Integrated DNA Technologies (IDT DNA, USA). The g-blocks were designed using SnapGene® by inserting the gene-of-interest protein sequences between the restriction enzyme (RE) sites in the pAF124 vector. The DNA sequences in the g-blocks were codon-optimized using IDT DNA’s codon-optimization tool. To facilitate Gibson Assembly, all g-blocks were designed with ∼25-30bp long regions of homology with the plasmid vector at the 5’ and 3’ terminal distal regions of the GOI’s. gbAF012 overlaps onto the EF1a promotor and encodes the signal peptide and mouse anti-6xHis (3D5) and overlaps onto the 5’ end of devil IGHG constant region.

#### 3.3.1 PCR amplify DNA fragments

1. The primers sha_IGHG.FOR and sha_IGHG.REV were designed with no extension to amplify the devil IGHG constant region (**Table 2**). PCR amplify the devil IGHG constant region from devil peripheral mononuclear cell RNA that has been converted to cDNA.
2. Set up and run a PCR reaction as previously described. Cycling conditions are as follows – 98 °C for 2 minutes, 35 cycles of 98 °C for 10 seconds, 62 °C for 15 seconds, and 72 °C for 45 seconds, followed by a final extension of 72 °C for 5 minutes.
3. Run products on a 1% agarose gel and purify the appropriate DNA fragments as previously described.

**Table 2.**
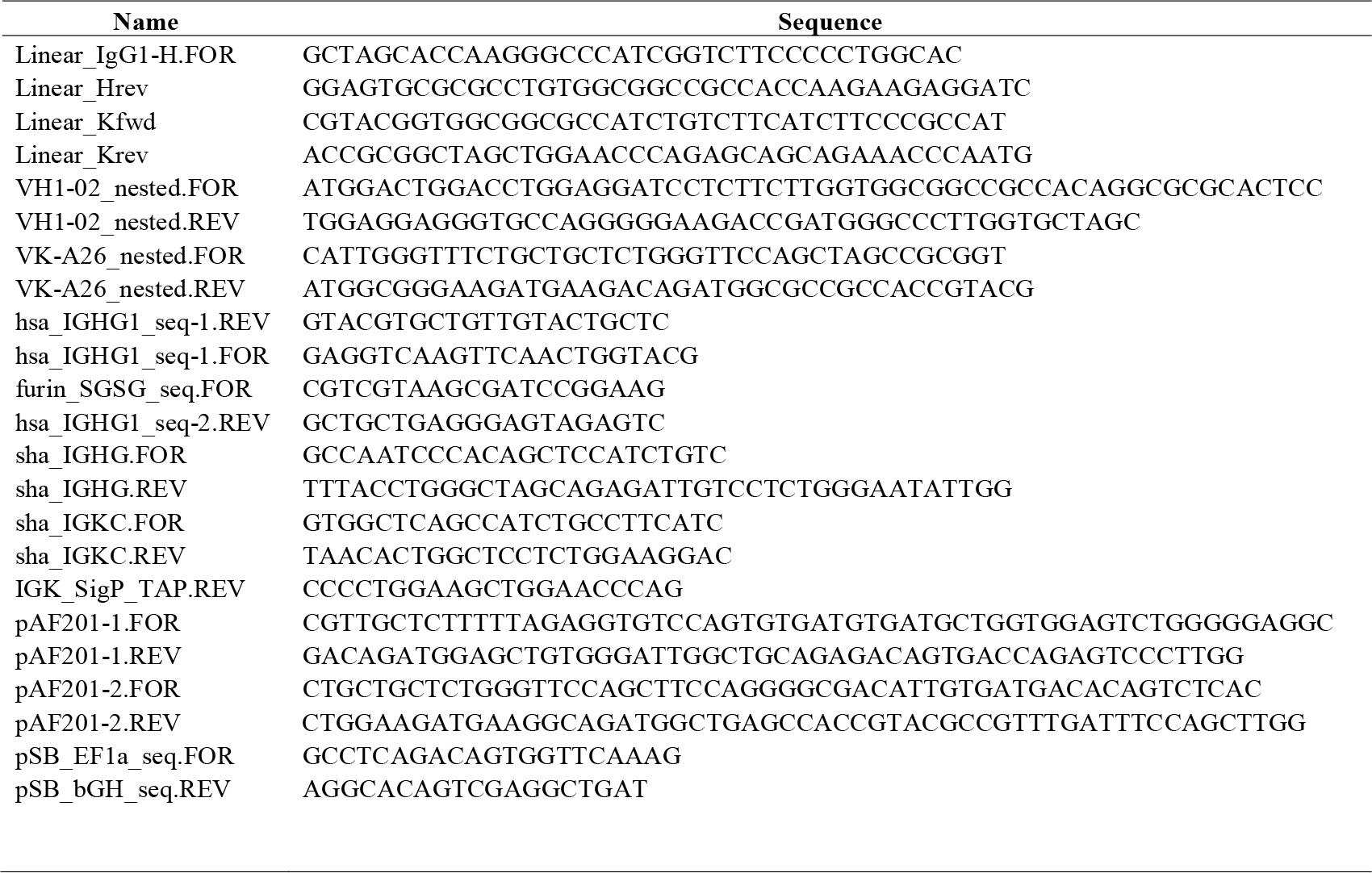
Primer for plasmid assembly and sequencing.

#### 3.3.2 Assembly of devil IgG anti-His (pAF200) expression vector

1. RE digest pAF124 with SfiI and assemble the devil IgG heavy and light chain constant region g-blocks and the linking fragment into the digested vector using Gibson assembly as described previously.
2. Sequence the assembled human IgG1anti-devil PDL1 expression vector.
3. Transfect the confirmed clone into CHO cells as described previously and collect supernatant.

#### 3.3.3 ELISA for detection of devil anti-His antibodies

1. Dilute recombinant Tasmanian devil interferon gamma (IFNγ) with a C-terminal 6xHis-tag to 1 μg/ mL in ELISA carbonate coating buffer (section 2.4.2). Add 100 μL into appropriate wells on a 96-well ELISA plate in triplicate. Negative control wells incubated with coating buffer only.
2. Seal the plate and incubate overnight at 4 °C.
3. Wash plate with PBS-T. Block plate with 100 μL of ELISA blocking buffer. Incubate for 1 hour at room temperature.
4. Wash plate 3 times with PBS-T. Add 100 μL of primary antibodies, recombinant human IgG1 anti-His or recombinant devil IgG anti-His transfected CHO cell supernatant to appropriate wells. Do not add any primary antibody to a subset of wells to serve as a negative control for primary antibodies. Incubate at room temperature for 1 hour (*see* **Note 18**).
5. Wash plate 3 times with PBS-T. Dilute a secondary antibody (mouse anti-Tasmanian devil IgG) 1:200 (v/v) in blocking buffer and add 100 μL to the devil anti-His wells. Incubate for 1 hour at room temperature.
6. Wash plate 3 times with PBS-T. Dilute tertiary antibodies anti-human IgG-HRP 1:20,000 and anti-mouse IgG-HRP 1:4000 in blocking buffer. Add 100 μL to appropriate wells (*see* **Note 19**). Incubate at room temperature for 30 minutes.
7. Wash plate 3 times with PBS-T. Add 100 μL of TMB peroxidase substrate to the wells. Plate can be read at 650 nm while colour is developing. Allow to develop until colour is obvious but then stop with 100 μL of 1 M hydrochloric acid.
8. Read using a Tecan Spark 20M plate reader at 450 nm (*see* **Note 20**).

#### 3.3.4 Assembly of tasmanian devil IgG anti-PDL1 (pAF201) expression vector

1. PCR amplify the IgG heavy chain linker fragment from the devil IgG anti-His vector (pAF200) backbone, and the anti-devil PDL1 heavy and light chain variable region fragments from the human IgG anti-devil PDL1 (pJD1) vector. as previously described. Primers pAF201-1.FOR and pAF201-1.REV, and pAF201-2.FOR and pAF201-2.REV were designed to amplify the anti-devil PDL1 heavy and light chain variable regions respectively, and primers sha_IGHG.FOR and IGK_SigP_TAP.REV were designed to amplify the linker fragment (**Table 2**).
2. Run samples on a 1% agarose gel and purify the amplified fragment as previously described (section 3.2.3).
3. RE digest the Tasmanian devil IgG anti-His expression vector pAF200 with EcoRV and BsiWI and assemble the mouse anti-devil PDL1 heavy and light chain variable region fragments amplified in section 3.4.1 and the linking fragment into the digested vector using Gibson assembly as described previously.
4. Sequence the assembled devilised IgG1 anti-PDL1 expression vector. Transfect the confirmed clone into CHO cells as previously described and collect supernatant.

#### 3.3.5 Test transfected CHO cell supernatant by flow cytometry to confirm antibody binding

1. Test the binding ability of the devil anti-PDL1 IgG antibody to devil PDL1 expressing CHO cells by flow cytometry as previously described with alteration after step 3.
2. Add 100 μL of neat Tasmanian devil anti-PDL1 IgG CHO cell supernatant to the cells and incubate for 30 minutes on ice. Wash twice with 100 μL of FACS buffer.
3. Add 100 μL of mouse anti-devil IgG [20] diluted 1:200 in FACS buffer. Incubate for 30 minutes on ice. Wash twice with 100 μL of FACS buffer.
4. Add 100 μL of anti-mouse IgG (Alexa Fluor™ 488) diluted 1:1000 in FACS buffer. Incubate for 30 minutes on ice. Wash twice with 100 μL of FACS buffer.
5. Resuspend cells in 400 μL DAPI in FACS buffer.
6. Analyse cells by flow cytometry along with controls.

## 4 Results

We tested the supernatant from CHO cells stably transfected with the pAF124 and pAF200 plasmids. The supernatant was tested in ELISAs for binding to a target protein with a C-terminal 6xHis tag. Binding was detected only when the target protein was present and the appropriate primary and secondary antibodies were included in the assay. Thus, the recombinant antibodies retain specificity for the C-terminal 6xHis tag.

The anti-PDL1 recombinant antibodies were tested using flow cytometry. CHO cell lines stably transfected with either devil CTLA4 or PDL1 were used as targets. Both the humanised and devilised anti-PDL1 antibodies retained their specificity for devil PDL1.

## 5 Discussion

This is the first report of a recombinant antibody with a Tasmanian devil heavy and light chain constant regions. Two different devil IgG chimeric antibodies retained binding to their target proteins. The devil IgG antibodies can be used *in vitro* for functional assays with devil immune cells to assess antibody-dependent cytotoxicity.

Both DFT1 and DFT2 cells have been shown to upregulate PDL1 in response to IFNγ treatment *in vitro*. Expression of cell surface PDL1 could be an important secondary immune evasion mechanisms for DFT1 and DFT2 cells. These chimeric antibodies could be used *in vivo* for anti-PDL1 immunotherapy more effectively than the original mouse IgG antibodies due to the mouse antibodies being viewed as fully foreign by the devil immune system. These chimeric antibodies retain the mouse variable regions, which could be targeted by the devil immune system if repeated doses of immunotherapies were administered. Fully devilised recombinant antibodies with devil variable regions would likely be most effective *in vivo*. This study takes a key step toward bringing immune checkpoint immunotherapy to wildlife disease.

## 6 Notes

1. The expression vector design is the key step of the process. All other steps will fail if the design is not effective.
2. These plasmids can also be used for transient transfections but the transposition is not needed for transient transfections. Transposition could reduce yield of the target protein compared to non-integrating plasmids.
3. Avoid UV light wavelengths as this can damage the DNA. Work quickly even when using blue light.
4. Overnight digest at ambient temperature works better for many restriction enzymes.
5. Loading 60 μL requires large wells. This can be accomplished by taping together two small comb tines to create a larger time and subsequently a larger well that can hold the required volume.
6. Use the NEBuilder® Positive Control pUC19 plasmid with an ampicillin resistance gene from the kit as a positive control reaction. Another reaction containing the plasmid vector and no inserts can be used as a negative control.
7. Thaw bacterial cells on ice just long enough to melt the ice. Do not thaw for longer than 10 minutes. Pre-chill sterile microfuge tubes on ice if transferring aliquots of bacteria to new tubes.
8. Use the bacteria as soon as they are thawed (∼5-10 minutes).
9. Avoid submerging the whole tube as this increases the chance of contamination.
10. Spread transformed bacteria evenly and gently over the LB agar plate, ensuring not to dig into the agar. Do this in a biosafety cabinet and semi-cover the plates as they dry to reduce the chance of contamination.
11. Serial dilutions of the bacteria can be used on different plates to ensure a single colony can be isolate. However, with an appropriate streaking pattern you can usually isolate individual colonies regardless of the bacterial dilution.
12. The plates containing the recombinant plasmid vector and positive control transformed cells should have a high number of colonies present, while there should not be any on the negative control plate. However, even a single colony on a plate can have the correctly assembled plasmid.
13. This will serve as the template for the PCR test and can be used to inoculate a broth culture downstream.
14. Using transfection grade plasmid purification kits may result in higher transfection efficiency downstream.
15. Monitor the cells daily. Massive cell death should occur within 1-3 days. Remove the floating cells every other day and replace with fresh, warm cRF5 with 800 μg/ mL hygromycin. Small colonies of cells should start forming after 2-5 days, allow these to reach ∼20 cells before passaging. Once all cells appear healthy in the flask the hygromycin dose can be lowered to 100-200 μg/mL. Expand and freeze vials of the cell for future use.
16. Ensure the cells have 2-5 days to secrete antibody into the supernatant after drug selection is complete before collecting the supernatant. The supernatant can be stored in the dark at 4 °C for up to 7 days post-harvest.
17. Use cell lines expressing another protein and/ or human antibodies against other targets as negative controls for flow analysis.
18. This antibody clone only binds His-tags at the C-terminus of the protein.
19. Use Anti-HIS-HRP on recombinant Tasmanian devil interferon gamma (IFNy) with 6xHis-tag as a positive control at a dilution of 1:2000.
20. Perform all ELISA conditions in triplicate.

## Acknowledgements

The authors would like to thank Alana de Luca, Sambavi Singarasa, and Amanda Patchett for helping with the laboratory research. The data for Figure 3 were generated in the Master’s thesis of Alana de Luca. This work was supported by ARC DECRA grant # DE180100484, University of Tasmania Foundation Dr. Eric Guiler Tasmanian Devil Research Grant through funds raised by the Save the Tasmanian Devil Appeal (2017, 2018), a Charitable organisation from the Principality of Liechtenstein, and a Select Foundation Fellowship to ASF.

**Fig. 3.**
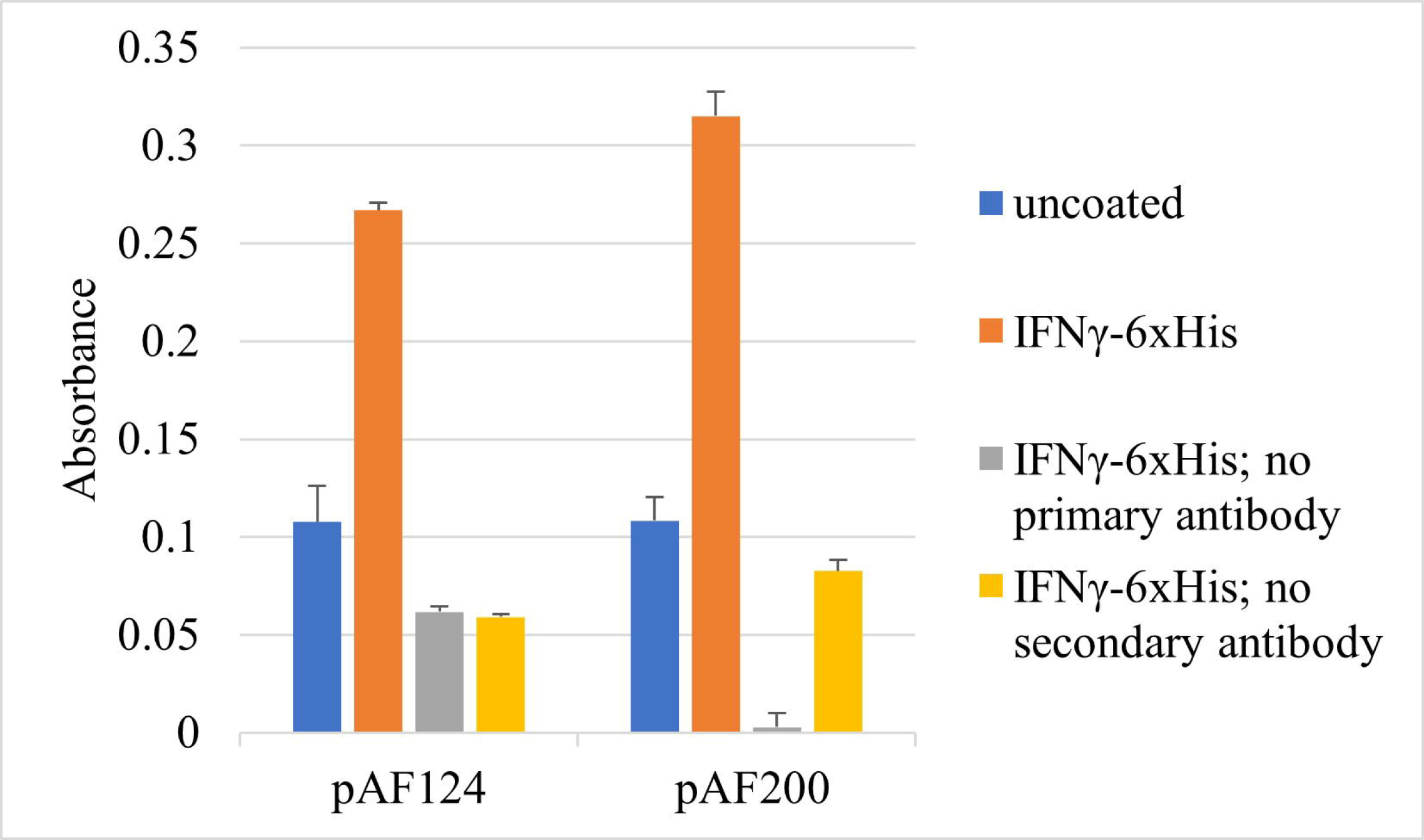
Chimeric anti-6xHis antibodies binds recombinant proteins with 6xHis in ELISAs. Humanised (pAF124) and devilised (pAF200) anti-6xHis binding to recombinant proteins with C-terminal 6xHis. Antibody combinations were performed in triplicate.

**Fig. 4.**
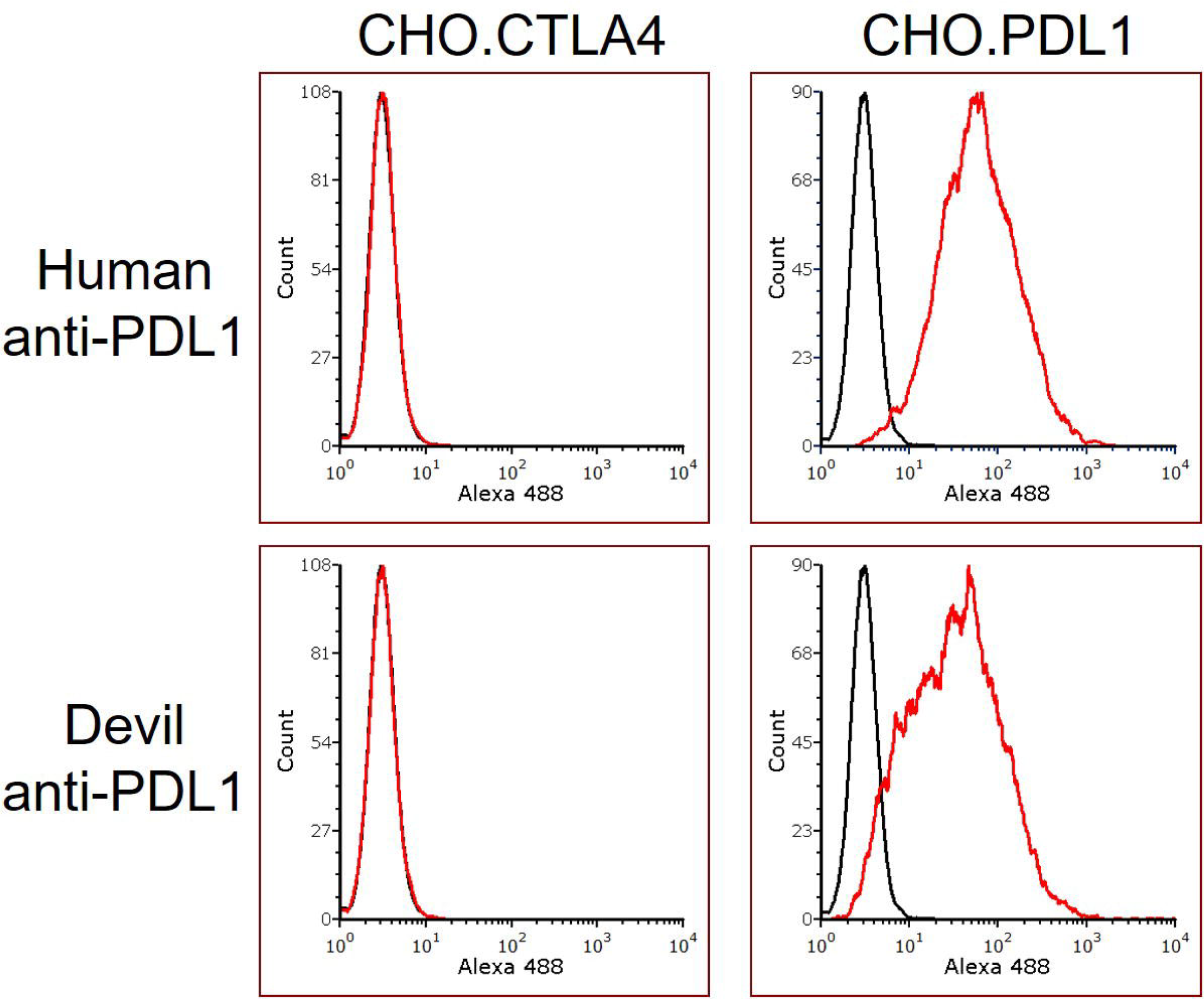
Chimeric anti-PDL1 antibodies bind to devil PDL1 in flow cytometry assays. CHO cells stably transfected with either devil CTLA4 or PDL1 were used as target cells. Recombinant anti-PDL1 antibodies bound to PDL1 transfected cells but not to CTLA4 cells.

